# Spike afterpotentials shape the in-vivo burst activity of principal cells in medial entorhinal cortex

**DOI:** 10.1101/841346

**Authors:** Dóra É. Csordás, Caroline Fischer, Johannes Nagele, Martin Stemmler, Andreas V.M. Herz

## Abstract

Principal neurons in rodent medial entorhinal cortex (MEC) generate high-frequency bursts during natural behavior. While *in vitro* studies point to potential mechanisms that could support such burst sequences, it remains unclear whether these mechanisms are effective under *in-vivo* conditions. In this study, we focused on the membrane-potential dynamics immediately following action potentials, as measured in whole-cell recordings from male mice running in virtual corridors (Domnisoru et al., 2013). These afterpotentials consisted either of a hyperpolarization, an extended ramp-like shoulder, or a depolarization reminiscent of depolarizing afterpotentials (DAPs) recorded *in vitro* in MEC stellate and pyramidal neurons. Next, we correlated the afterpotentials with the cells’ propensity to fire bursts. All DAP cells with known location resided in Layer II, generated bursts, and their inter-spike intervals (ISIs) were typically between five and fifteen milliseconds. The ISI distributions of Layer-II cells without DAPs peaked sharply at around four milliseconds and varied only minimally across that group. This dichotomy in burst behavior is explained by cell-group-specific DAP dynamics. The same two groups of bursting neurons also emerged when we clustered extracellular spike-train autocorrelations measured in real two-dimensional arenas (Latuske et al., 2015). No difference in the spatial coding properties of the grid cells across all three groups was discernible. Layer III neurons were only sparsely bursting and had no DAPs. As various mechanisms for modulating the ion-channels underlying DAPs exist, our results suggest that the temporal features of MEC activity can be altered while maintaining the cells’ spatial tuning characteristics.

**Significance Statement:** Depolarizing afterpotentials (DAPs) are frequently observed in principal neurons from slice preparations of rodent medial entorhinal cortex (MEC), but their functional role *in vivo* is unknown. Analyzing whole-cell data from mice running on virtual tracks, we show that DAPs do occur during behavior. Cells with prominent DAPs are found in Layer II; their inter-spike intervals reflect DAP time-scales. In contrast, neither the rarely bursting cells in Layer III, nor the high-frequency bursters in Layer II, have a DAP. Extracellular recordings from mice exploring real two-dimensional arenas demonstrate that grid cells within these three groups have rather similar spatial coding properties. We conclude that DAPs shape the temporal but not the spatial response characteristics of principal neurons in MEC.

**Author contributions:** All authors designed research. DÉC, CF, and JN performed research and analyzed data (equal contribution). AVMH wrote and edited the paper with support from MS and the other authors.

## Introduction

Principal neurons in the superficial layers of medial entorhinal cortex (MEC) show rich temporal behavior, from slow depolarization ramps (Domnisoru et al., 2013, Schmidt-Hieber and Häusser, 2013), spike locking and phase precession in the theta band (Hafting et al., 2008; Reifenstein et al., 2012), to gamma-band activity (Chrobak and Buzsáki, 1998; Colgin et al., 2009) and burst sequences with instantaneous firing rates of up to 300 Hz (Latuske et al., 2015). The spatial firing fields of one particular MEC cell class, namely grid cells, form hexagonal lattices spanning the explored two-dimensional environment (Hafting et al. 2005). Notably, not every grid cell participates to the same degree in these temporal phenomena. In particular, there are two sub-classes of grid cells, those that burst frequently and those that do not or only rarely generate bursts (Mizuseki et al., 2009, Latuske et al., 2015, Ebbesen et al., 2016). But what is the mechanism behind the MEC bursts and what role do they play for spatially selective neurons, such as grid cells?

In this study, we tested the hypothesis that burst activity of principal neurons in MEC is shaped by cell-intrinsic membrane-potential dynamics. Two mechanisms come to mind. Bursts could reflect action-potentials (APs) riding on high-frequency membrane-potential oscillations, akin to theta-band coupling in MEC Layer-II stellate cells (Alonso and Klink, 1993; Engel et al. 2008; see also Hasselmo, 2013; Newman and Hasselmo, 2014). Alternatively, bursts could result from AP-triggered membrane-potential dynamics that increase the probability of further discharges.

Indeed, slice experiments have shown that depolarizing afterpotentials (DAPs) arise in a majority of principle cells in superficial MEC layers (Alonso and Klink, 1993; Canto and Witter, 2012). DAPs are at the center of triphasic deflections following an AP, sandwiched between fast and medium after-hyperpolarization (fAHP and mAHP). The DAP maximum occurs some five-to-ten milliseconds after the AP and peaks a few millivolts above the fAHP minimum. In stellate cells, DAPs become more pronounced when neurons are hyperpolarized, whereas the reverse is true for pyramidal neurons (Alessi et al., 2016). Not all cell types associated with grid-like rate maps have DAPs in vitro, however. In particular, layer-III neurons are reported to have no DAPs (Canto and Witter, 2012).

DAPs do not only agree in their relevant time scale with intra-burst inter-spike intervals (ISIs); in vitro, DAPs also play a causal role for bursting. Alessi et al. (2016) reported that during DAPs the AP current threshold was reduced such that the cells’ average excitability increased by over 40%. Conversely, neurons without strong DAPs did not burst at the beginning of an AP train (Canto and Witter, 2012).

To test the functional relevance of DAPs under in-vivo conditions, we analyzed whole-cell recordings from mice moving on a virtual linear track (Domnisoru et al., 2013) and could show that DAPs play a decisive role for burst firing in MEC Layer-II neurons: Cells with DAPs were bursty and their intra-burst ISIs were compatible with the DAP mechanism. ISI distributions of the other Layer-II cells were highly uniform and had a sharp peak at 4.1±0.2 ms (SD across this cell group). The remaining neurons were sparsely bursting and those with known location resided in Layer III, apart from one pyramidal cell in Layer II. The results are compatible with our findings for extracellular recordings from open-field arenas (Latuske et al., 2015). In addition, bursty cells with and without DAP did not differ in their spatial coding properties. As the ion-channels underlying DAPs can be modulated in many ways, our analysis suggests that temporal features of grid-cell activity can be altered to serve different functions without affecting the cells’ spatial tuning characteristics.

## Materials and Methods

### Data

We analyzed data from two separate studies in navigating wild-type (C57BL/6) male mice. Data set “D” (Domnisoru et al., 2013) contained voltage traces from whole-cell recordings sampled at 20 kHz in head-fixed animals. These mice ran on cylindrical treadmills through virtual corridors. Data set “L” (Latuske et al., 2015) contained tetrode data (Sampling frequency: 20 or 24 kHz) obtained during movements in a real square arena (70 × 70 cm).

### Cell selection

Data set D: The original data set contained recordings from 51 cells of which 27 had been classified as grid cells by Domnisoru et al. (2013). One grid-cell recording was partially corrupted and excluded. Two grid cells had mean firing rates above 10 Hz and were removed to allow for an unbiased comparison with data set L, which contained only cells with firing rates below 10 Hz to exclude interneurons. From the 24 neurons that had been classified as non-grid cells two cells had firing rates above 10 Hz and the action potentials of six other cells did not meet our criteria (see below under “Membrane-potential dynamics”). This resulted in 40 neurons from data set D, namely 24 grid cells and 16 non-grid cells.

Data set L: After removing cells for which the animal trajectories showed artifacts, 522 principal cells were identified using the same criterium (mean firing rate < 10 Hz) as in Latuske et al. (2015). Out of those cells, 115 cells had been classified as grid cells by these authors. Ten of the 522 cells were not considered further as their inter-spike interval (ISI) distributions differed strongly from all other cells in that they had not a single inter-spike interval below 8 ms. Similarly, to avoid artifacts in the cluster analysis of the cells’ spike-time autocorrelations, seven cells with sparse autocorrelations were removed (see below for details). Altogether, this led to 505 cells in data set L, 112 grid cells and 393 non-grid cells.

### Spike-train characterization

The firing rate of a cell was defined as number of spikes divided by the total duration of the recording. For the graphical illustrations, spike-time autocorrelations and ISI distributions were calculated from binned data (bin width: 1 ms). To compute the peak location and width of ISI distributions, the recorded time difference between each pair of successive spikes was represented by a Gaussian kernel with a standard deviation of 1ms, centered at the mesaured time difference. These individual kernel density (KD) estimates were summed up across the entire recording. The analogous procedure was used for autocorrelations.

The location of the ISI peak was determined as the inter-spike interval for which the KD estimate was maximal. Similarly, the width of the ISI distribution was defined as full width at half maximum. The mean ISI and its standard deviation were calculated from all ISIs, the coefficient of variation (CV) was defined as the ratio between standard deviation and mean.

A burst was defined as a sequence of at least two spikes with ISIs shorter than 8 ms. The fraction of ISIs smaller than 8 ms was calculated relative to all ISIs and serves as a measure for the cell’s burstiness. An event is a burst or an isolated spike. The fraction of single spikes was defined as the number of spikes that do not belong to a burst divided by the number of all events.

### Principal-component (PC) analysis

For both data sets, autocorrelations were calculated for time lags between 0 ms and τ_max_ = 50 ms. To weigh all neurons equally autocorrelations were normalized to unit area. Principal components were calculated after binning (bin width: 1 ms). To reduce spurious effects caused by sparse normalized autocorrelations, cells with more than 75% empty bins (the maximum value for data set D) were removed (one grid cell and six non-grid cells in data set L). For the same reason, 5 (2) cells of data set D (data set L) that had relatively few spikes (< 130) were excluded when PC components were calculated but are included in the further analysis. To test the robustness of the PC analysis of the D data, the maximal time lag τ_max_ was varied between 30 ms and 100 ms (see also Results).

### Identification of neuron classes

For the D data set, visual inspection of the two-dimensional space spanned by the first two PCs suggested two main cell groups, whose arrangement was determined by k-means clustering with k=2 clusters. To test the robustness of the k-means clustering for the L data set, cluster analyses were performed on the 50-dimensional binned autocorrelations as well as in PC spaces with N=2-4 dimensions. The clustering quality was estimated through silhouette scores (Rousseeuw et a., 1987).

### Membrane-potential dynamics

The whole-cell voltage traces contained sizeable fluctuations that reflected synaptic inputs and potential movement artefacts. To obtain reliable information about the membrane potential before and after an action potential, AP-triggered averaging was performed. The APs themselves varied in amplitude and width, both within and across the different recordings, suggesting that the recording quality varied in time; the slowly decaying AP amplitudes and increasing width of some cells indicated run-down effects. To guarantee a good recording quality and to obtain reliable estimates of the subthreshold membrane-potential dynamics on the time scales relevant for fAHPs and DAPs, we focused on well isolated APs (no further APs within 25 ms before and after the trigger AP), and required the individual AP amplitudes to be larger than 40 mV (measured relative to the membrane potential 10 ms before the AP maximum) and APs width to be smaller than 1 ms.

The pre-AP voltage slope was calculated from the cell’s average AP-triggered voltage trace within the last 10 ms before AP onset; AP onset was determined by a threshold crossing (15 mV/ms) in the average AP-triggered voltage trace.

For cells with DAPs, the fAHP amplitude ΔV_fAHP_ was defined as the average voltage minimum during the fAHP relative to the voltage at AP onset. This means that ΔV_fAHP_ is negative for DAP cells (see Fig. 1). The DAP-deflection ΔV_DAP_ was defined as the difference between the voltage level at the DAP peak and at the minimum of the preceding fAHP. It is positive for cells with DAPs. The time interval between the AP peak and the following fAHP minimum is denoted by Δt_fAHP_, the time interval between the AP peak and the following DAP maximum is called Δt_DAP_.

**Figure 1.**
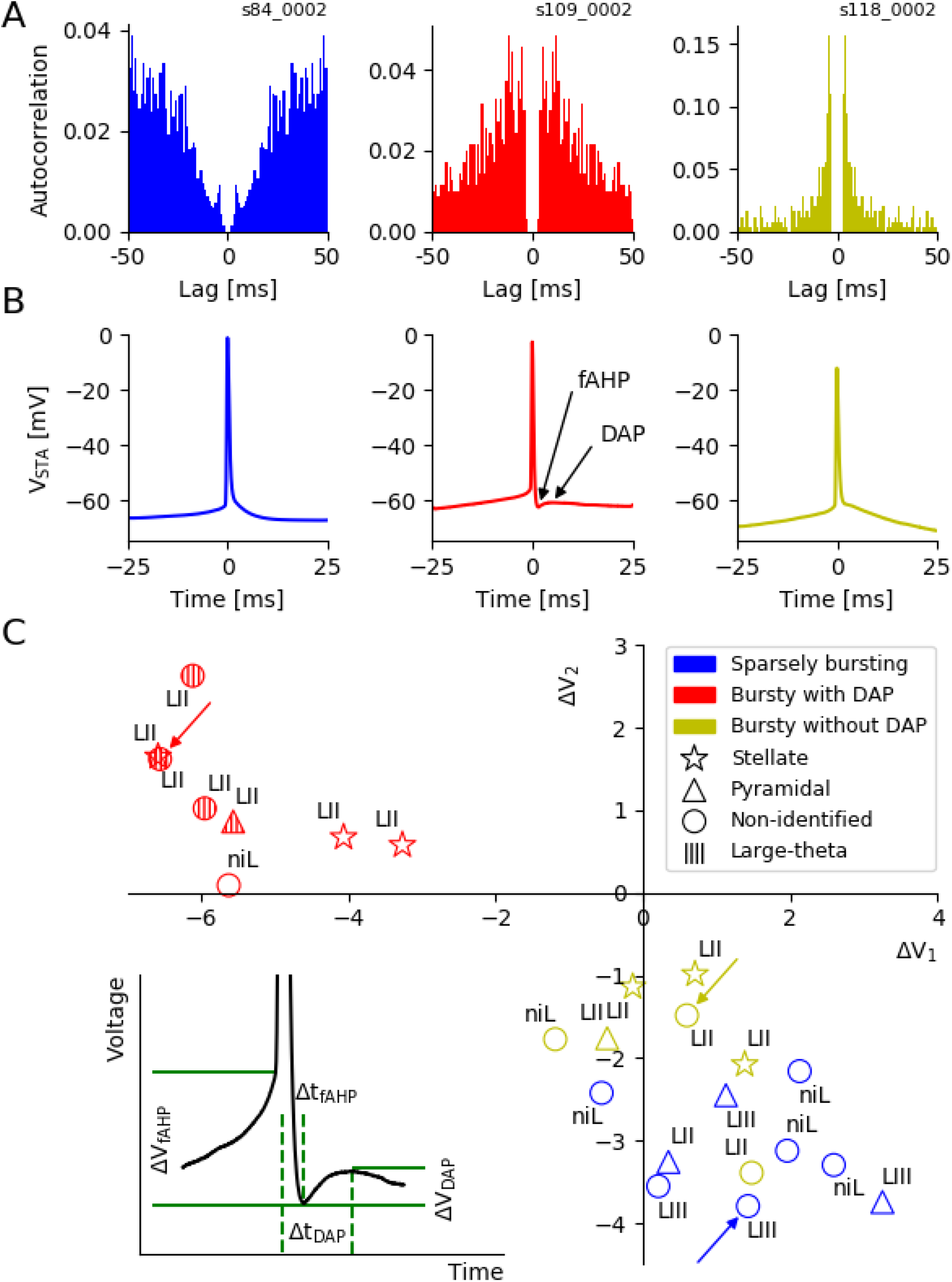
Spike afterpotentials of MEC grid cells from mice moving in virtual corridors. **A**, typical examples of grid-cell burst behavior. Left panel: autocorrelation function of a sparsely bursting cell; middle panel: a bursting cell with broad autocorrelation flanks; right panel: a bursting cell with sharply peaked autocorrelation. Note the different scale. **B**, grid cells differ in their spike afterpotentials. Left panel: a monotone repolarization that is gradually slowing down; middle panel: fast hyperpolarization (fAHP) followed by a depolarizing afterpotential (DAP); right panel: a short repolarization that abruptly turns into a much slower voltage decay, which may include an initial flat shoulder. **C**, characterization of spike afterpotentials. Inset: Definition of parameters. Main panel: group data. To characterize afterpotentials for cells without DAPs, the two parameters ΔV_1_ and ΔV_2_ take the role of ΔV_fAHP_ and ΔV_DAP_. The new parameters are determined in the same way as ΔV_fAHP_ and ΔV_DAP_, at times Δt_1_ and Δt_2_, respectively. These times are obtained from averages of Δt_fAHP_ and Δt_DAP_ across the population of DAP cells. Finally, for DAP cells, ΔV_1_ and ΔV_2_ are used instead of ΔV_fAHP_ and ΔV_DAP_, to simplify the notation.

To compare the afterpotentials of all neurons studied, the definitions of ΔV_fAHP_ and ΔV_DAP_ had to be generalized to cells without DAP. This was done (a) for grid cells only, and (b) for all neurons. To this end, we calculated the population averages 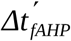 and 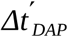 across the respective DAP cells (grid cells: n=8, all neurons: n=15). We then used these mean time intervals (grid cells: 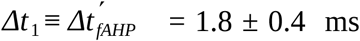, 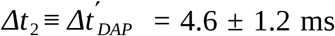, all neurons: 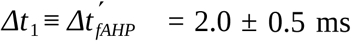, 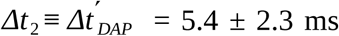) to determine voltage changes corresponding to ΔV_fAHP_ and ΔV_DAP_ for cells without DAP. These are called ΔV_1_ and ΔV_2_, respectively. For notational simplicity, we will use these terms for grid cells, too, but here, they denote the ΔV_fAHP_ and ΔV_DAP_ values measured at the cell-specific Δt_fAHP_ and Δt_DAP_ values.

### Spatial coding properties

For data set L, grid score and head-direction score were calculated as in Sargolini et al. (2006), the spatial information as in Skaggs et al. (1996).

### Experimental design and statistical analysis

We analyzed data recorded by Domnisoru et al. (2013) and Latuske et al. (2015) and refer the reader to these two publications for details on the experimental design. All our analyses were performed in Python 2.7.6. Specific statistical tests used are stated throughout the text. The Kruskal-Wallis test, the Kolmogorov-Smirnov test, the Chi-square test and the median tests are taken from scipy.stats. The linear regression, the principal component analysis and the k-means clustering are taken from scikit-learn.

### Bootstrapping

To assess the fAHP and DAP parameters, we bootstrapped the AP-triggered voltage traces of a cell by using sampling with replacement and repeated this procedure 10000 times to obtain mean values and standard errors.

## Results

The temporal firing characteristics of principal neurons in the medial entorhinal cortex (MEC) of behaving rodents vary strongly from cell to cell, even if their mean firing rates are almost identical (Latuske et al., 2015). Some neurons rarely fire with inter-spike intervals shorter than 8 ms; their spike-time autocorrelations have a pronounced dip at short time lags (Fig. 1*A*, left). Other cells show an autocorrelation peak in the 5-15 ms range with broad flanks (Fig. 1*A*, middle) and yet other cells have autocorrelations that are sharply peaked at even shorter lags (Fig. 1*A*, right). Not distinguishing between the second and third group of neurons, these cells have been termed “bursty” by Latuske et al. (2015) whereas the first group has been called “non-bursty” by these authors. Since “non-bursty” neurons generate bursts from time to time, too, we will call them “sparsely bursting”, in line with Simmonet and Brecht (2019).

We wondered whether differences in the *in vivo* spike patterns of bursty neurons could be explained at a mechanistic level by differences in their intrinsic single-cell dynamics and whether – for the spatially tuned cells within the total population – differences in the cells’ temporal discharge patterns were reflected in their spatial tuning properties. To this end, we analyzed whole-cell recordings from mice moving on a linear track in virtual reality (Domnisoru et al., 2013) and extracellular recordings from mice navigating in two-dimensional environments (Latuske et al., 2015). We start with an analysis of grid cells and then extend our approach to non-grid cells.

### Grid cells differ in the voltage deflections following an action potential

We first focus on the intracellular linear-track data as these provide information about both spike times and membrane-potential dynamics.

The time courses of the membrane potentials recorded by Domnisoru et al. (2013) show striking cell-to-cell differences within the first ten milliseconds following an action potential (AP). Three distinct types of behavior can be distinguished from the spike-triggered voltage traces:

i. a monotone repolarization that gradually slows down (Fig. 1*B*, left panel)
ii. a fast afterhyperpolarization (fAHP) followed by a depolarizing afterpotential (DAP), as shown in Fig. 1*B*, middle panel,
iii. a short repolarizing phase that abruptly turns into a much slower voltage decay, which may include a flat shoulder (Fig. 1*B*, right panel).

To quantify these distinct behaviors, we used parameters that capture two salient features of cells exhibiting DAPs – the voltage minimum during the fAHP and the voltage peak during the DAP (see the lower left inset in Fig. 1*C*). The “fAHP-depth” ΔV_fAHP_ measures the voltage minimum relative to the membrane potential at AP onset. This minimum occurs at some time Δt_fAHP_ after the AP peak. The “DAP-deflection” ΔV_DAP_ measures the difference between the membrane potential at the DAP peak and the fAHP minimum. The DAP peak is attained at some time Δt_DAP_ after the AP peak (Fig. 1*C*).

To extend the ΔV_fAHP_ and ΔV_DAP_ measures to voltage traces of cells with no detectable DAP, two time intervals corresponding to Δt_fAHP_ and Δt_DAP_ need to be defined. For concreteness, we used the population means across all grid cells with a DAP (n=8), resulting in 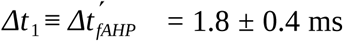 and 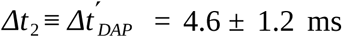. We then determined the voltage differences corresponding to ΔV_fAHP_ and ΔV_DAP_ at these two time points, and named them ΔV_1_ and ΔV_2_, respectively. For DAP cells, we define ΔV_1_ = ΔV_fAHP_ and ΔV_2_ = ΔV_DAP_ to simplify the notation. These settings mean that cells with a shoulder or slow voltage decay (Fig. 1*B*, right panel) have zero or small ΔV_2_ irrespective of their ΔV_1_ value whereas large negative ΔV_2_ values indicate a strong decline in membrane potential until around five milliseconds after the action potential.

Within the parameter space spanned by ΔV_1_ and ΔV_2_, neurons fall into two distinct groups (Fig. 1*C*) – cells with a pronounced DAP (negative ΔV_1_ and positive ΔV_2_) and cells with no detectable DAP (negative ΔV_2_), which typically have also no fAHP (positive ΔV_1_). Results from a bootstrapping analysis (see Materials and Methods) underscore the reliability of the DAP and fAHP parameter estimates (see Fig. 1-1).

All measured neurons that had a DAP and whose location was known, resided in Layer II. Five of these eight cells had large theta-band membrane potential oscillations and have been called “large theta cells” by Domnisoru et al. (2013). Cells without a DAP (n=16) were located in both layers II and III.

### Grid cells differ in their spike-train characteristics

To capture the diversity of spike discharge patterns we carried out a principal component analysis on the spike-time autocorrelations of the 24 intracellularly recorded grid cells (Fig. 2*A*), as has been done for extracellular recordings (Latuske et al, 2015). We restricted our attention to autocorrelations on short time-scales, in particular to the region between 0 and 50 ms after a spike. We found that the first two principal components, PC1 and PC2, explain 66% and 18% of the cell-to-cell variability, respectively, whereas the contribution from PC3 adds only another 4%. Together, PC1 and PC2 thus account for 84% of the variability. This value changes by less than 3% when τ_max_ is varied between 30 ms and 100 ms (data not shown) and starts to decrease for shorter or longer maximal lag. These findings indicate that a two-dimensional PC representation of the grid-cell autocorrelations in the 0-50 ms range describes the essence of the cell-to-cell variability.

**Figure 2.**
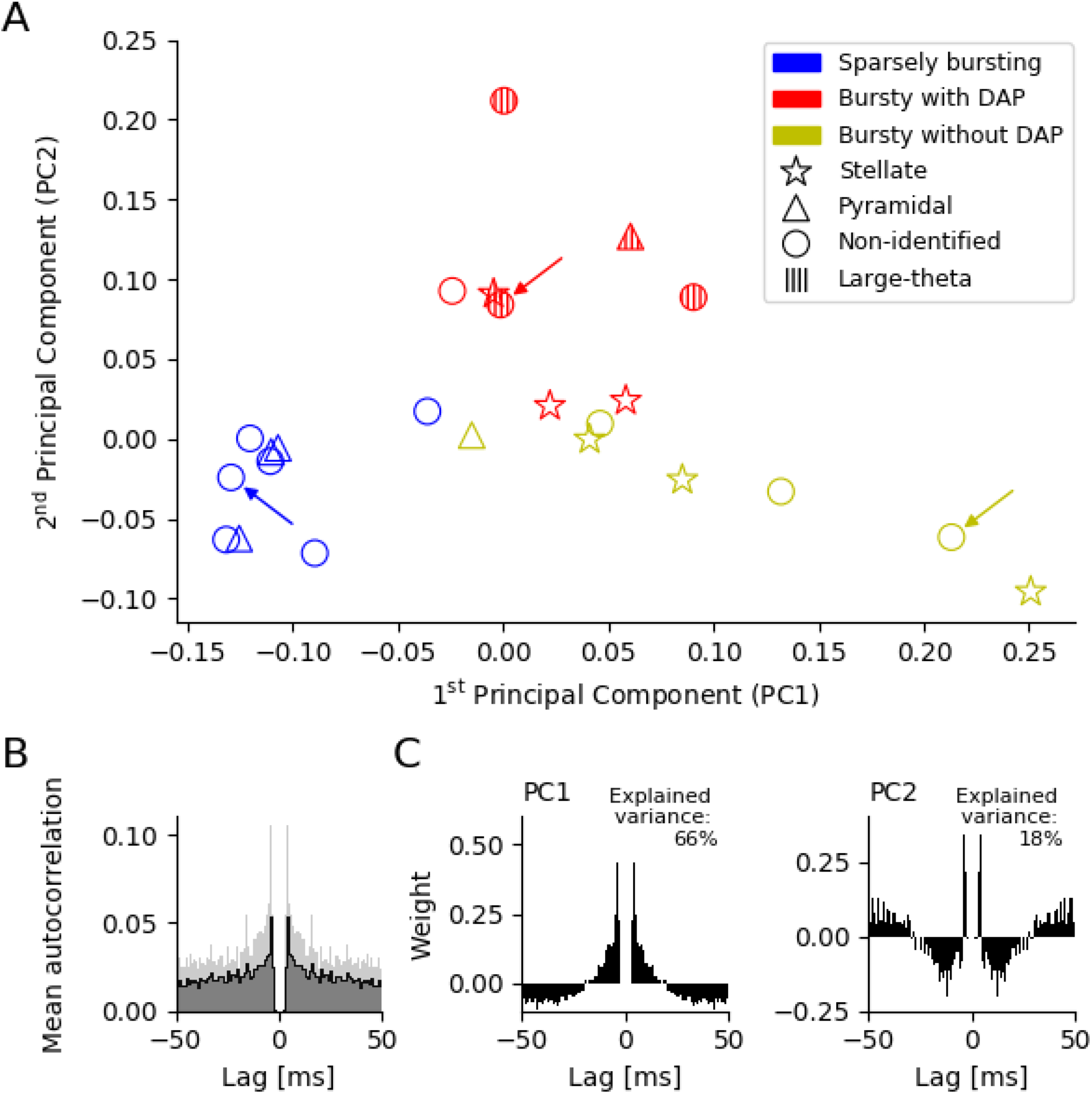
Spike-time autocorrelations of MEC grid cells from mice moving in virtual corridors. **A**, visual inspection of spike-time autocorrelations suggests a separation in two major groups, sparsely bursting and bursting. Based on the intracellular measurements (see Fig.1), the group of bursting neurons can be subdivided in cells with DAP (“ BD^ᴑ^ ”), which are shown in red, and cells without detectable DAP (“ BD^−^ ”), shown in yellow. The arrows mark the example neurons shown in Fig.1. One cell has intermediate properties and is assigned to the sparsely-bursting group (blue) by k-means clustering (k=2). **B**, mean autocorrelation. **C**, the first two principal components of the spike-time autocorrelations. The pronounced peaks in B and C demonstrate that inter-spike intervals of around 4 ms are indicative of both the mean grid-cell discharge patterns and their cell-to-cell variability.

The mean autocorrelation is highly peaked at a lag τ of around 4 ms (Fig. 2*B*), and so are both principal components (Fig. 2*C*). This indicates that brief activity bursts in the 250 Hz range play an important role for both the mean grid-cell discharge patterns and their cell-to-cell variability.

Within this two-dimensional representation (Fig. 2*A*), neurons without a DAP, shown in blue and yellow, have a negative or only small positive second principal component and strongly vary in their first principal component. Cells with large negative PC1 are sparsely bursting as the example in the left panel of Fig. 1*A*, whose position in the PC1/PC2 space is marked with a blue arrow in Fig. 2*A*. Cells with positive PC1 burst like the cell in the right panel of Fig. 1*A* (yellow arrow in Fig. 2*A*). Cells with a DAP have positive PC2, only a small PC1, and are also bursting, though with a much broader peak in their autocorrelation as demonstrated by the example in middle panel in Fig. 1*A*, marked with a red arrow in Fig. 2*A*.

This grouping is based on visual inspection of the AC principal components and might not properly distinguish between bursting and sparsely-bursting neurons with small PC1. To better discriminate between these two cell groups, we carried out a k-means clustering with k=2. The analysis suggested that 9 cells should be classified as sparsely bursting (“SB”) neurons; based on their intracellular characteristics, the remaining 15 bursting (“B”) cells are either DAP cells (“ BD^ᴑ^ ”) or cells without detectable DAP (“ BD^−^ ”). The same clusters emerge if the spike data from the first and second half of each experiment are treated separately (data not shown), and provide evidence for the robustness of our approach.

All bursty neurons whose anatomical position was classified by Domnisoru et al. are located in Layer II, none in Layer III (two bursty cells were not assigned to a layer). Furthermore, unlike suggested by reports from rats (Ebbesen et al., 2016), bursty neurons are more likely to be stellate than pyramidal cells (6 versus 2 cells), in agreement with the larger abundance of stellate cells compared to pyramidal cells (Alonso and Klink, 1993). There was no detectable difference in the morphology of bursty neurons with and without DAP: Within the BD^−^group (n=8), there were three stellate cells, one pyramidal neuron and four non-identified cells. Within the BD^−^group (n=7), there were three stellate cells, one pyramidal neuron and three non-identified cells. In contrast, not a single non-bursty cell was identified as a stellate cell (pyramidal and non-identified cells: 3 and 6 out of n=9, respectively) and non-bursty neurons may have a tendency to reside in Layer III (three cells versus one cell in Layer II; 5 cells were not classified). Finally, in the ΔV_1_–ΔV_2_ representation (Fig. 1*C*), BD^−^neurons overlap with SB cells, but tend to have less negative ΔV_2_ values and the ΔV_1_ and ΔV_2_ values within the group of bursty neurons are correlated with a slope of −0.49 (standard error: 0.06). The three groupings are robust, as confirmed by bootstrapping and indicated by error bars in Fig. 1-1.

### Post-AP dynamics explain the spike-train characteristics of bursty grid cells

After dividing the cells into three groups based on their spike-train autocorrelations and DAP characteristics, we compared the group averages for the grid cells’ intracellular voltage traces (Fig. 3). Confirming the impression from individual cells, sparsely-bursting neurons show a smooth and monotone AP down-stroke (Fig. 3*A*, left panel), bursty cells with DAP exhibit a local voltage minimum followed by repolarization (Fig. 3*A*, middle panel), and compared to SB cells, bursty cells without DAP tend to have two phases of repolarization: an initial AP downstroke followed abruptly by a slower rate of repolarization, yielding a kink in the voltage traces (Fig. 3*A*, right panel).

**Figure 3.**
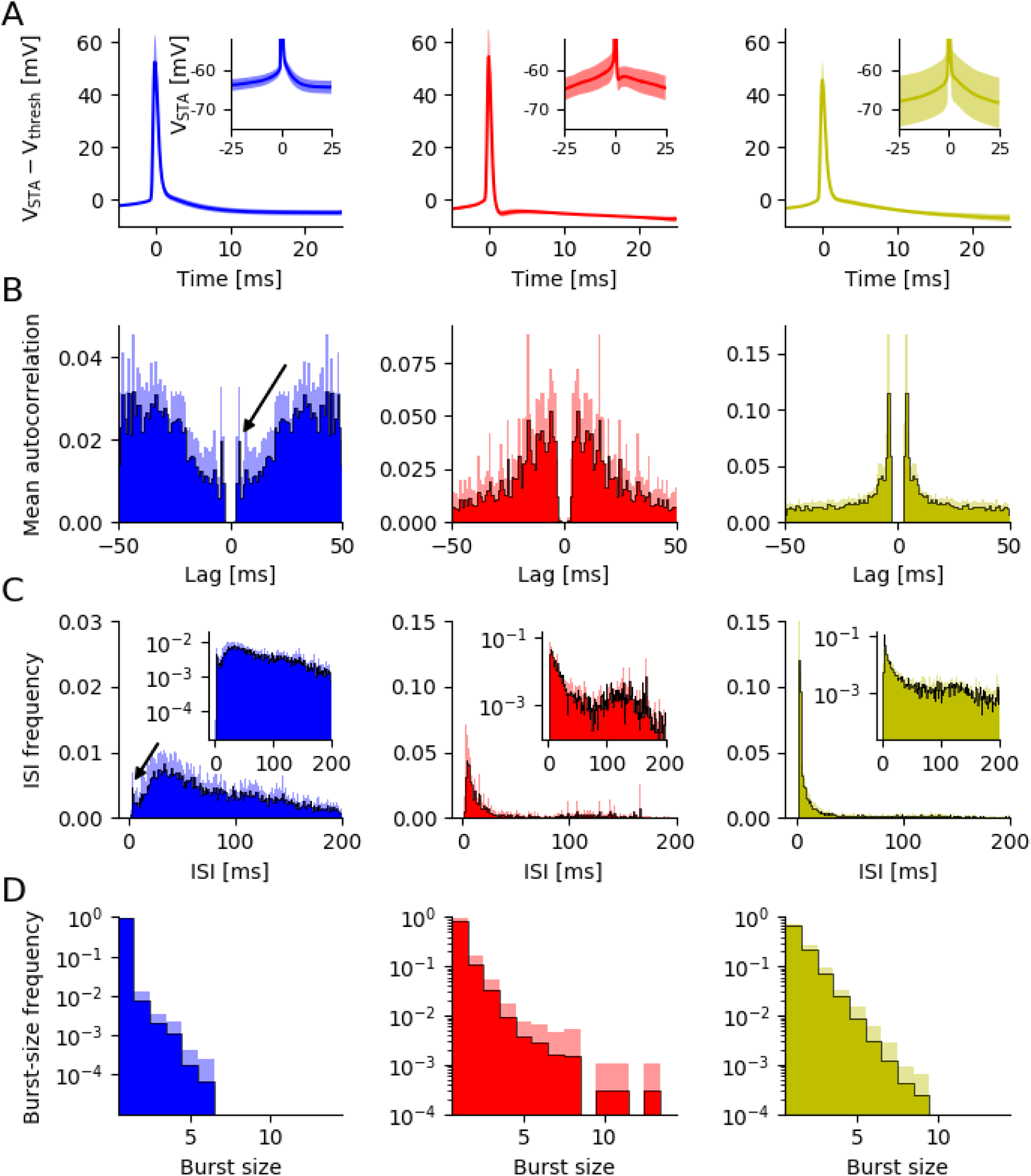
Group-level analysis of MEC grid cells from mice moving in virtual corridors. **A**, population average of the spike-triggered membrane potential for isolated action potentials (no further AP within 25ms before and after the AP). The main plot shows data that were aligned to AP onset before the group average was taken, the inset illustrates the absolute membrane potential values. Shaded area: standard deviation **B**, population-averaged autocorrelation functions. **C**, population-averaged inter-spike intervals distributions. Arrows in B and C highlight bursts of sparsely bursting cells, inset in C with logarithmic time scale emphasizes theta-band activity. **D**, population-averaged intra-burst spike count distributions.

To visualize these distinct features, the spike-triggered voltage traces were averaged for each neuron and then aligned to the cell’s mean voltage at AP onset. Without voltage alignment (see insets), slight differences in membrane potential at AP onset are apparent (SB: −59.42 ± 1.22 mV, BD^ᴑ^: −57.12 ± 3.09 mV, BD^−^: −61.40 ± 6.83 mV) but not significant (p(BD^ᴑ^, BD^−^)= 0.25; p(BD^ᴑ^, SB)=0.15; p(BD^−^, SB)=0.96, Kruskal Wallis), and a larger variability of the afterpotentials within the two groups of bursty neurons. The voltage slope during the last 10 ms before AP onset does not differ significantly between BD^ᴑ^ and BD^−^neurons (BD^ᴑ^: 0.50 ± 0.03 mV/ms; BD^−^: 0.48 ± 0.08mV/ms; p=0.35, Kruskal Wallis) but does so when sparsely-bursting and bursty neurons are compared (B: 0.49 ± 0.06 mV/ms; SB: 0.34 ± 0.05 mV/ms; p=0.00015, Kruskal Wallis). Finally, visual inspection suggests that there are no high-frequency membrane-potential oscillations, which would have been expected if the bursts resulted from electric resonances.

The averaged autocorrelations (Fig. 3*B*, left panel) and inter-spike intervals (Fig. 3*C*, left panel) of sparsely-bursting cells show that although these neurons rarely generate spike sequences with short ISIs – only 2% of all their ISIs are less than 8ms – if they do fire such bursts, there is a pronounced short ISI that is only 4.30 ± 0.81ms long (see the black arrows in Figs. 3B and C). Both types of bursty neurons exhibit prominent ISI- and autocorrelation peaks at short time scales (Fig. 3B, C, middle and right panels). Population averages within each group show that the most likely ISI of cells without a DAP is significantly shorter than that of cells with a DAP (4.12 ± 0.12 ms vs. 6.96 ± 3.73 ms, p=0.01, Kruskal Wallis); the same is true for the autocorrelation peaks (4.13 ± 0.11 ms vs. 9.46 ± 4.41 ms, p=0.001, Kruskal Wallis). These differences are readily explained by the different time courses of the post-spike voltage deflections: The rapid fAHP time course of BD^ᴑ^ cells strongly reduces the chance that a second AP is fired directly after the first AP, whereas in BD^−^ cells the down-stroke of the first AP stops abruptly at depolarized levels, often above the AP threshold (see Fig. 1*B*, 3A), resulting in a rather short absolute refractory period (mean of 10 % shortest ISI in BD^−^ cells: 3.45 ms versus 5.24 ms in BD^ᴑ^ cells). Consistent with this picture, the DAP opens a wide “window of opportunity” for a second AP in BD^ᴑ^ cells, resulting in broader ISI distributions (6.7 ± 3.27 ms vs. 3.66 ± 0.36 ms, p=0.01, Kruskal Wallis) for BD^ᴑ^ cells, compared to BD^−^ cells. Finally, a direct role of the post-AP dynamics in burst behavior is also suggested by the observation that for BD^ᴑ^ cells, the most likely ISI mirrors Δt_DAP_, the time interval between an AP and the succeeding DAP peak (no difference of median values, p=0.61, median=5.13 ms, median test).

The intrinsic voltage dynamics alone do not explain why sparsely-bursting cells have much broader ISI distributions (see Fig. 3*C*). Other features correlated with sparsely bursting behavior, though. All identified SB cells were pyramidal neurons, whereas only one out of four BD^−^ cells were pyramidal; four of the five SB cells were in LIII, whereas BD^−^ cells were solely found in LII. By contrast, the BD^−^ and BD^ᴑ^ groups were not distinguished by cell-type or the layer in which they were found, which makes it unlikely that anatomical differences can explain the observed variations in the spike trains of bursty neurons.

*In vitro*, stellate cells often produce spike doublets or brief bursts (Alessi et al., 2016). We tested whether spike trains *in vivo* showed similar preferences. For this purpose, we computed the frequency with which cells fired two, three, or more spikes in a burst. The frequency of bursts with exactly n spikes decreases monotonically with n, and does so for all three cell groups. Doublets and triplets were not overrepresented (Fig. 3*D*). In 21 out of the 24 cells, the distribution for n is consistent with an exponential distribution (Chi-Square Goodness of Fit Test for the correlation between linear fit and data after logarithmic transformation; p>0.05). There is thus no preferred burst size or “unit of information”, such as a spike doublet or triplet. In the spirit of hippocampal “complex spike bursts” (Ranck, 1973), a grid-cell burst can be regarded as just a sequence of two or more spikes with short inter-spike intervals. The exact choice of the cutoff threshold is not critical; qualitatively similar results were obtained using ISI thresholds from 8 ms up to 15 ms (data not shown).

### Spike-train characteristics of bursty cells are conserved across 1D and 2D environments

So far, the analysis was based on a relatively small number of neurons recorded in head-fixed animals running in a virtual linear corridor (Domnisoru et al., 2013). To explore whether these data generalize to other experimental conditions, we analyzed a complementary data set that contained 112 grid cells from mice that foraged at random in a square environment (Latuske et al., 2015). Although these extracellular recordings do not offer direct access to the membrane-potential dynamics, they might still reveal signatures of the different post-AP dynamics. In particular, we expected that grid cells would not only show the bursty vs. non-bursty dichotomy described by Latuske at al. (2015), but that there would also be qualitative differences within the bursty subpopulation.

To facilitate the comparison between the two data sets we kept the maximum lag of 50 ms and the 1 ms-binning when analyzing spike-time autocorrelations. To minimize observer bias, k-means cluster analyses were performed on the 50-dimensional raw autocorrelations as well as in principal-component spaces with N=2-4 dimensions. We analyzed the robustness of the k-means clustering results by calculating silhouette scores for each value of k (Rousseuw et al., 1987). Irrespective of the dimensionality N of the data, separation into three clusters led to the best performance; the clusters hardly changed when using different numbers of principal components. Cluster assignment was stable, regardless of whether the autocorrelations were computed from the first or second half of a cell’s spike train: 91.1% of the cells kept their cluster identity (Fig. 4-1) from the first to the second half when using N=3 principal components. To compare these data with the previous results, we again plot the first two principal components against each other (Fig. 4*A*). The mean autocorrelation (Fig. 4*B*) closely resembled the one obtained from the virtual-track data (see Fig. 2*B*); and so did the principal components (cf. Fig. 4*C* and Fig. 2*C*), as quantified by their similarity (the scalar product computed for time lags up to 50 ms) of 0.86 for PC1 and 0.63 for PC2 when the two experimental conditions are compared.

**Figure 4.**
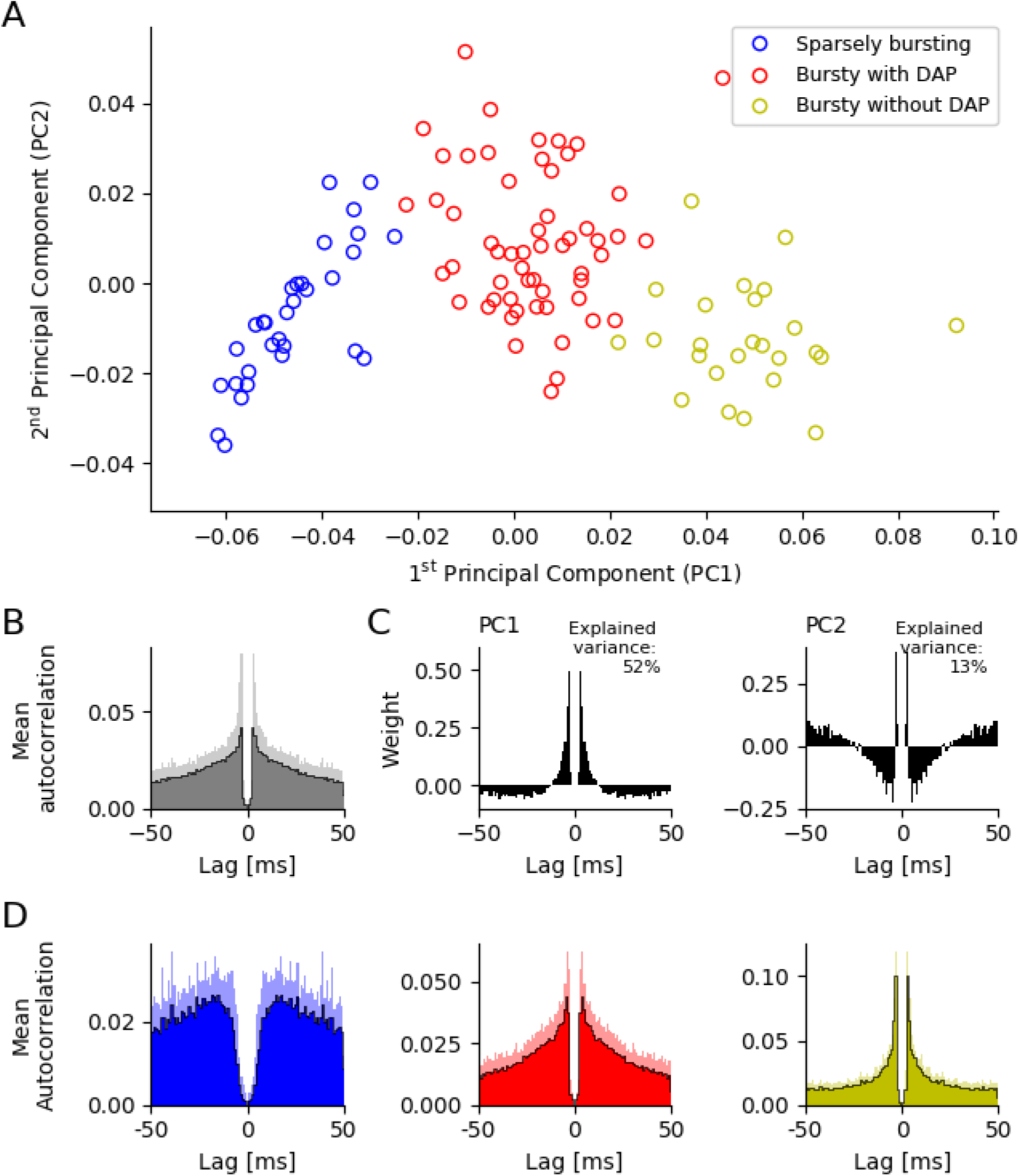
Spike-time autocorrelations of MEC grid cells from mice moving in open arenas. **A**, k-means cluster analysis (k=3) of spike-time autocorrelations. **B**, mean autocorrelation across the grid cells. **C**, the first two principal components of the spike-time autocorrelations. The sharp peaks in B and C again demonstrate the prevalence of short inter-spike intervals (here around 3.5 ms) in the mean grid-cell discharge patterns as well as their cell-to-cell variability. **D**, autocorrelations averaged across all neurons from each cell group reveal a striking similarity between cells recorded on virtual tracks and in open fields. Strongest deviations are shown by non-bursting/sparsely bursting cells in the Latuske et al. vs. Domnisoru et al. data.

There is also a high similarity between the corresponding group-averaged autocorrelations in both data sets (SB cells: 0.93, BD^−^ cells: 0.93, BD^ᴑ^:cells: 094), cf. Fig. 4*D* and Fig. 3*B*. This is remarkable as the cluster analysis of the open-field data (Latuske et al., 2015) only reflects the overall structure of the grid-cell autocorrelations and is not informed by intracellular measurements. There is, however, one prominent difference between both data sets. The sparsely bursting neurons recorded by Latuske et al. (Fig. 4*D*, left panel) fire hardly any spike within the first few milliseconds (so that the authors named them “non-bursty” neurons) and their autocorrelation has a pronounced peak at around 15 ms. The autocorrelation function of the sparsely bursting neurons recorded by Domnisoru et al. (Fig. 3*B*, left panel) exhibits a local peak at around 4 ms, and grows more slowly, with a local maximum at 30-40 ms. On the other hand, the average autocorrelations of the BD^ᴑ^ and BD^−^ cells are almost identical when compared across both experimental conditions.

In the virtual-track data, the autocorrelations of the BD^−^neurons (n=7) peak at 4.13 ms with a cell-to-cell variability of 0.11 ms (SD). The autocorrelations of the corresponding cells from the open-field recordings (n=25) have their peak at 3.56 ms (SD: 0.27 ms). There are biophysical differences between intra- and extracellular signals that might contribute to a longer delay being measured for the intervals between successive spikes (Anastassiou et al., 2015). The two experimental conditions also differ in the measured grid-field sizes: these are larger in virtual reality than open-field environments (c.f., Domnisoru et al., 2013, supp. Fig 9). Consistent with this observation, other spike-train measures also reflect longer timescales in virtual reality versus open fields, e.g., the most likely ISI (Fig. 5*A*), the width of the ISI histogram (Fig. 5*B*) or the fraction of inter-spike-intervals below 8 ms (Fig. 5*C*). Within each experimental setting, however, the three cell groups exhibited the same temporal features, albeit on slightly different timescales. On the other hand, the firing rates (Fig. 5*D*) are rather similar across experimental conditions and cell groups, which might reflect a general network-level regulation of the average firing rate.

**Figure 5.**
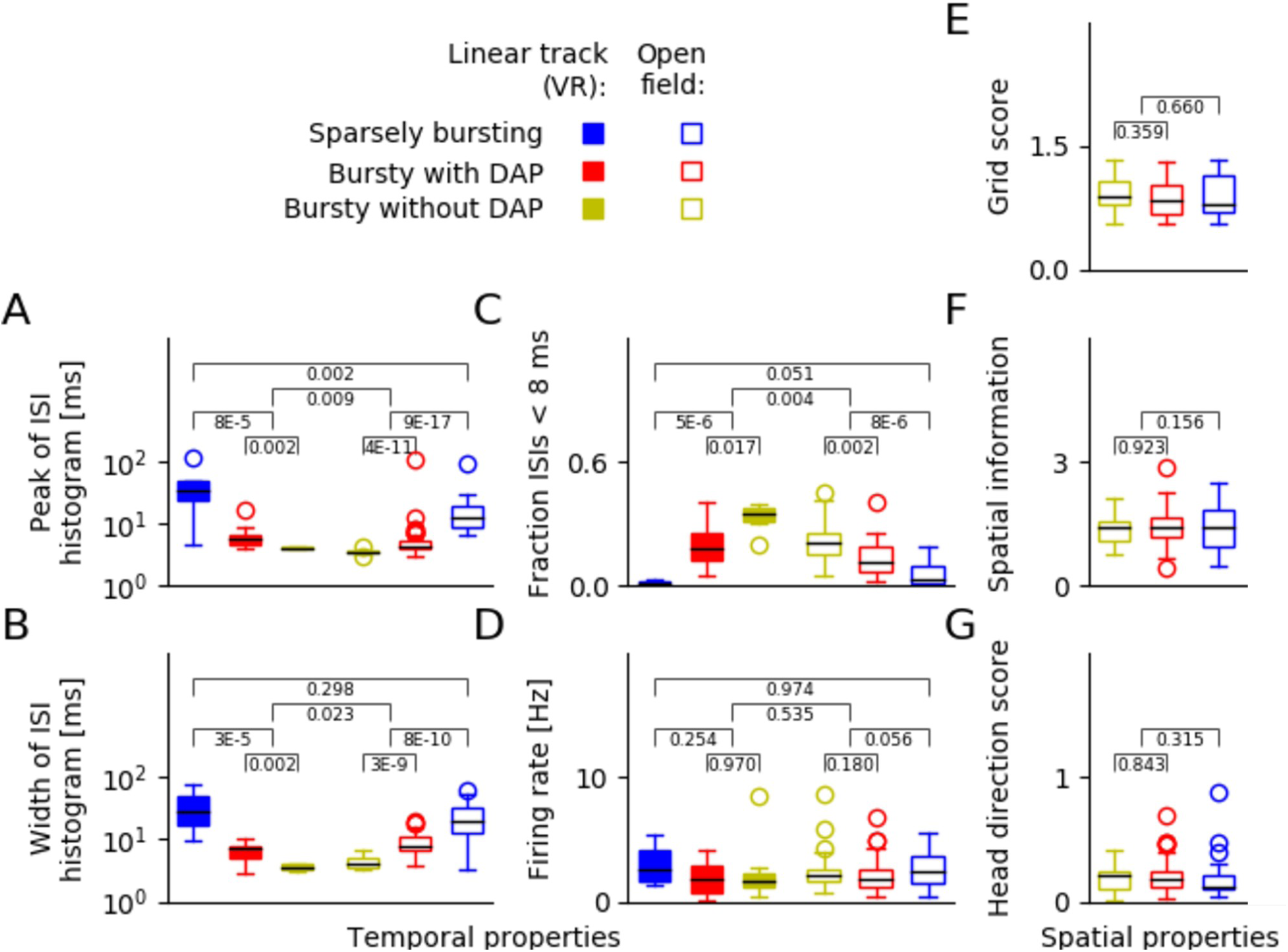
Grid-cell spike-train characteristics across data sets and spatial coding. **A-D**, comparison of linear-track data (filled symbols) and data from the open arena (un-filled symbols); **E-G**, spatial properties of grid fields recorded in the open arena. **A**, ISI peak, i.e., most likely inter-spike interval. **B**, width of the ISI distribution. **C**, fraction of ISIs below 8ms (“burstiness”). **D**, mean firing rates. **E**, grid score. **F**, spatial information. **G**, head direction score. Despite strong differences in temporal spike-train characteristics (**A-C**), mean firing rates (**D**) and spatial coding properties (**E-G**) of grid cells are largely conserved across all three cell groups.

### Spatial response properties are shared across all three grid-cell groups

In the next step of our analysis, we asked whether the pronounced differences between the temporal response characteristics of the three grid-cell groups translate into differences in their spatial firing patterns. The study of Latuske et al. (2015) had shown that this was *not* the case when one compares bursty with sparsely-bursting grid cells. However, the two groups of bursty neurons might still differ in their spatial behavior. To obtain reliable field estimates, we used the open-field data for this analysis. We tested various measures, including grid score (Sargolini et al., 2006), spatial information (Skaggs et al., 1996), and head-direction score (Sargolini et al., 2006), but could not detect any significant differences between BD^ᴑ^and BD^−^ cells (Fig. 5E-G).

Slice experiments show that depolarizing afterpotentials of stellate cells can be modulated; if the holding potential is decreased, the amplitude ΔV_DAP_ of the following DAP increases, and it decreases whenever the holding potential is increased (Alessi et al., 2016). As demonstrated by Domnisoru et al. (2013), a grid cell is depolarized when the animal is located in a firing field of that cell and hyperpolarized in the out-of-field regions. We therefore wondered whether a BD^ᴑ^ cell might preferentially generate DAP-mediated bursts when one of its grid field is entered, as the membrane-potential ramp might facilitate larger DAPs and thus make DAP-mediated burst firing in these neurons more likely.

To test this hypothesis, we took open-field data from Latuske et al. (2015) and investigated in detail whether spikes belonging to the bursts of a BD^ᴑ^ cell had an above-chance probability to occur at the edges of its firing fields and, more generally, whether those spikes differed in their spatial statistics from other spikes of the same neuron, in the spirit of a place-cell study by Harris et al. (2001). In particular, we analyzed the distribution of spike distances from the respective firing-field centers as well as topological features of the discharge patterns of BD^ᴑ^ cells, with special focus on inter-spike intervals expected for DAP-triggered bursts. Despite extensive efforts, we could not find any significant differences between the spatial firing characteristics of BD^ᴑ^ versus BD^−^ cells. As a complementary check, we used the data from Domnisoru et al. (2013) to search whether DAP deflections measured in the firing fields of BD^ᴑ^ cells were smaller than the DAP deflections of out-of-field spikes but did not find any obvious changes either.

These findings suggest that despite the striking differences in the spike-train patterns of bursty cells with and without DAP, these differences have no obvious consequences for the cells’ spatial tuning properties. Temporal variations in the membrane potential, in particular the large theta-oscillations observed in some bursty grid cells, are uncorrelated with the animal’s trajectory and may easily mask less prominent spatial dependencies. In fact, decoupling of spatial and temporal tuning characteristics might endow the system with added plasticity and computational flexibility.

### Bursty grid cells: One continuum or two clusters?

Since BD^ᴑ^ and BD^−^ cells showed indistinguishable spatial tuning, we reconsidered their partition into two distinct groups based on their temporal firing characteristics. Could it be that the data are better described as a single group with continuously varying parameters?

To answer this question, we went back to the grid-cell data from Domnisoru et al. (2013) and analyzed how the cells’ salient spike-train characteristics depended on the two biophysical parameters ΔV_1_ and ΔV_2_ (Fig. 6). The mean firing rate does not correlate with ΔV_1_ and ΔV_2_ (Fig. 6*A*,B). There are also no significant dependencies within the BD^ᴑ^ and BD^−^groups (ΔV_1_: p_BD+_=0.88, p_BD-_=0.24; ΔV_2_: p_BD+_=0.58, p_BD-_=0.90, as tested by independently shuffling the two coordinates of the respective data points and computing the Pearson correlation for each new sample. The p-value is given by the fraction of samples for which the correlation value was larger than in the original sample.).

**Figure 6.**
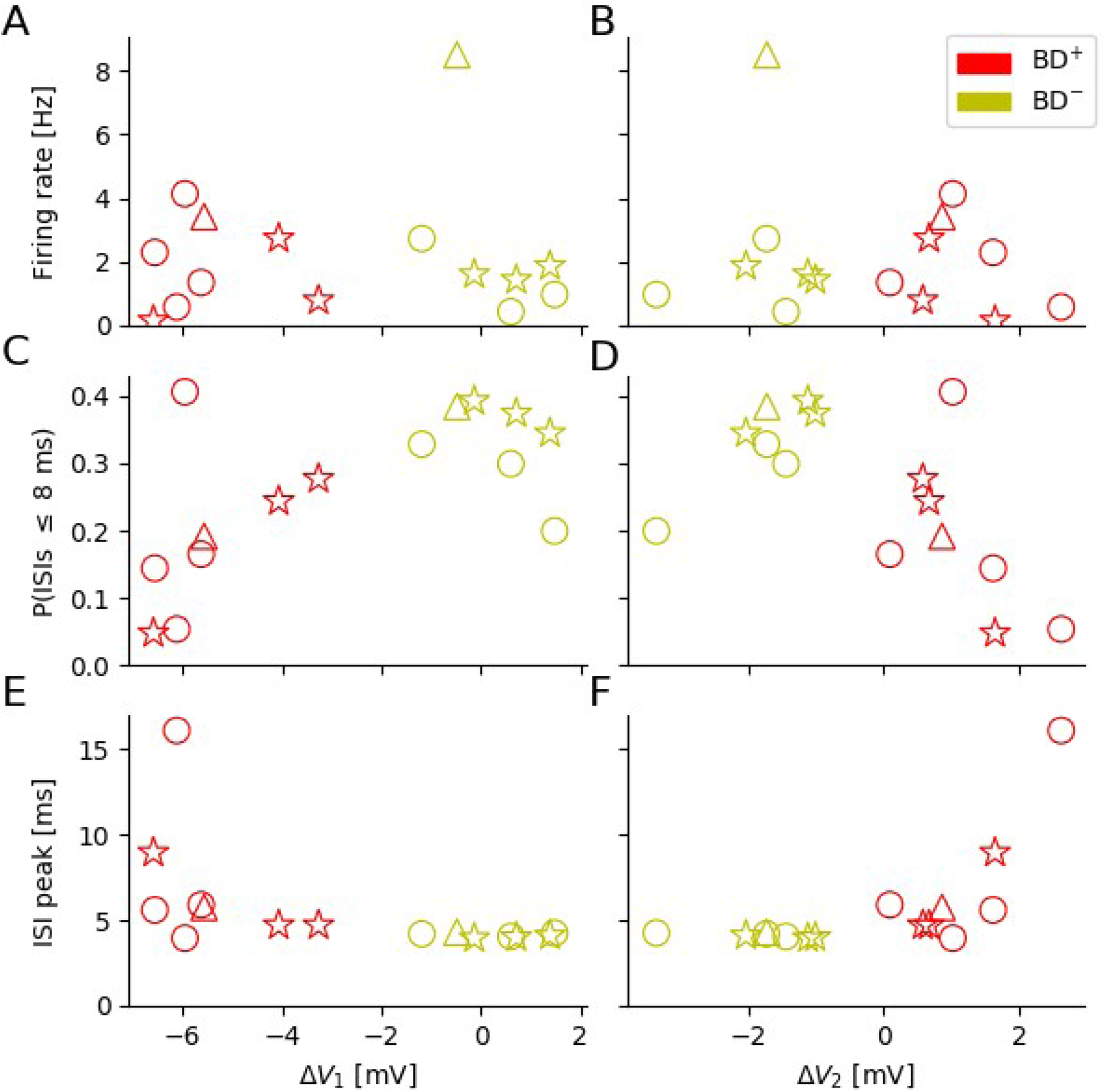
Cluster structure of the bursty-grid-cell population. Dependence of key spike-train parameters on the amplitude of fast afterhyperpolarization (ΔV_1_) and afterdepolarization (ΔV_2_). **A** and **B**, mean firing rates. **C** and **D**, fraction of ISIs below 8 ms (“burstiness”). Across the entire population of bursty neurons, the larger ΔV_1_, the more frequent are short ISIs **E** and **F**, location of ISI peak. While the firing rates do not exhibit a trend, neither within the two cell groups nor across the groups, the other quantities depicted show trends that differ from the null-hypothesis (no increase/decrease as a function of ΔV_1_ or ΔV_2_). The data also suggest that the population of bursty neurons either forms one joint though under-sampled cloud or contains two distinct sub-populations. In either case the spike-train characteristics do depend on the cells’ DAP properties.

A different picture emerges when the fraction of short ISIs (below 8 ms) is considered (Fig. 6*C,D*). Visual inspection suggests a joint trend for BD^ᴑ^ and BD^−^ cells; the larger ΔV_1_, the more frequent are short ISIs (Fig. 6*C*). With a p-value of 0.013, this trend is statistically significantly different from the null-hypothesis (no increase/decrease as a function of ΔV_1_), and in agreement with our earlier functional interpretation of depolarizing afterpotentials: For negative ΔV_1_ (i.e., BD^ᴑ^ cells), cells quickly hyperpolarize, making very short ISIs rare.

Consistent with this observation, the location of the ISI peak tends to grow for increasingly negative ΔV_1_ (Fig. 6*E*) and increasingly positive ΔV_2_ (Fig. 6*F*) if the entire population of bursty neurons is considered (ΔV_1_: p=0.04, ΔV_2_: p=0.01). Within the BD^−^ population, however, the ISI peak does hardly vary at all, as emphasized before.

These results indicate that there is no clear-cut answer to the question whether the population of bursty neurons forms one joint though under-sampled cloud or contains two distinct sub-populations. More importantly, however, is the observation that in either case, certain spike-train characteristics do depend on the cells’ individual DAP properties, which supports the view that DAPs do not only exist under in-vivo conditions but may also play a functional rule.

### Non-grid cells show same DAP and spike-train characteristics as grid cells

In the last step of our analysis, we asked whether non-grid cells differed from grid cells in their DAP behavior or spike-time autocorrelation characteristics. To this end, we first determined the DAP parameters ΔV_1_ and ΔV_2_ for all 40 neurons on linear tracks in virtual reality. As shown in Fig. 7*A*, non-grid cells (represented by “x”) fall into the same data clouds as the grid cells (represented by “o”) when these two intracellular measures are considered. Similarly, a principal-component analysis of the spike-train autocorrelations of the entire data set (Fig. 7*B*) exhibits a two-dimensional structure that is highly reminiscent of that when only grid cells are taken into account (c.f., Fig. 2*A*). The same is true for the principal-component analysis of the spike-train autocorrelations of the open-field data from Latuske et al. (2015) shown in (Fig. 7*C*). In fact, even the principal components themselves are highly similar when computed for grid cells only or for non-grid cells only (Fig. 7-1). Moreover, the entire data set exhibits the same ambiguity concerning the “one continuum vs. two clusters” question (Fig. 6-1) as when only grid cells are considered (Fig. 6).

**Figure 7.**
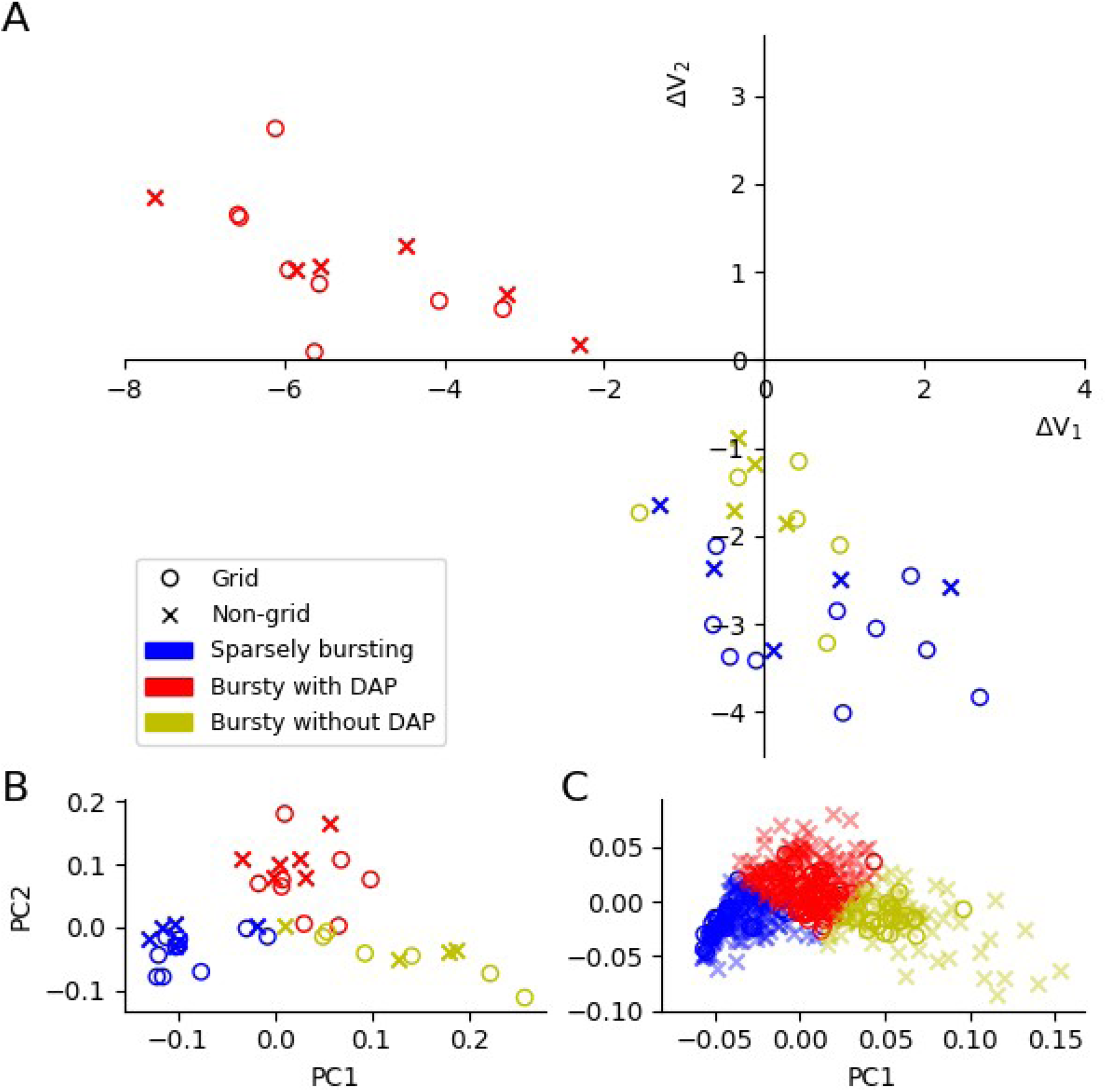
Comparison of the characteristics of grid cells and non-grid cells. **A**, quantification of spike afterpotentials as in Fig. 1C but now for all principal cells recorded on the virtual linear track. **B**, PC analysis of spike-time autocorrelations as in Fig. 2A but now for all principal cells recorded on the virtual linear track. **C**, PC analysis of spike-time autocorrelations as in Fig. 4A but now for all principal cells recorded in the open environment. The high similarity between grid cells and non-grid cells suggests that the three functional cell classes identified in this study are universal across all principal cells in the superficial MEC layers.

Consistent with this observation, the remarkably small cell-to-cell variability in the autocorrelation peaks of BD^−^ grid cells (virtual linear track: 4.13 ± 0.11ms; open field: 3.56 ± 0.27ms) is also shared by the non-grid BD^−^ cells (virtual linear track: 4.05 ± 0.25 ms, open field: 3.67 ± 0.60 ms), and these peaks are significantly shorter than the autocorrelation peaks of the non-grid BD^ᴑ^ cells (virtual linear track: 9.29 ± 3.53 ms, p=0.02, Kruskal Wallis; open field: 6.99 ± 3.52ms, p=1.15e-29, Kruskal Wallis). All data: mean values ± standard deviation.

Taken together, our findings demonstrate not only that the DAP characteristics of grid cells have no consequences for their spatial firing properties (c.f., Fig. 5E-G) but that in addition, non-grid cells and grid cells fall into the same three subgroups – sparsely bursting neurons, bursty with DAP and bursty without DAP. Both results suggest that DAPs and burst firing are not critical for spatial navigation.

## Discussion

Tetrode recordings in freely moving rats (Mizuseki et al., 2009, Ebbesen et al., 2016) and mice (Latuske et al. (2015) have shown that principal neurons in superficial MEC layers come in two functional subclasses, cells that burst frequently and others that do not or only rarely burst. Our analysis of whole-cell data from mice running on linear virtual tracks (Domnisoru et al., 2013) suggests that principal cells located in Layer-III tend to be “sparsely bursting” (SB) and that they do not generate depolarizing afterpotentials (DAPs), in agreement with previous slice studies in rats (Canto and Witter, 2012).

Bursty neurons varied strongly in the overall shape of their autocorrelations (as shown for grid cells in Fig. 3*B*) and their inter-spike interval (ISI) distributions (Fig. 3*C*). This diversity can be understood in terms of the cell-specific shapes of spike afterpotentials: Neurons without a DAP (“BD^−^ cells”) had inter-spike interval (ISI) distributions that peaked sharply at around four milliseconds and varied only minimally across that group of cells whereas the ISIs of neurons with a DAP (“BD^ᴑ^ cells”) were most frequent between 5 and 15 ms.

At first sight, the gap between BD^ᴑ^ and BD^−^ cells in the ΔV_1_-ΔV_2_ diagram (Fig. 1*C*) speaks against a continuum of bursty grid cells and rather points to the existence of two separate subgroups. This impression might, however, be due to a sampling artefact; there are only 15 such cells with intracellular recordings in the data set from Domnisoru et al. (2013). We therefore investigated the dependencies of various spike-train characteristics on ΔV_1_ and ΔV_2_ (Fig. 6). The smooth behavior of some measures, such as the burstiness, i.e., the fraction of ISIs below 8ms, or the ISI-peak location, and the lack of any sharp transitions in the other measures, support the assumption of one single, though sparsely populated group of neurons. Although based on small numbers, the equal stellate-to-pyramidal-cell ratio (3:1) of the BD^ᴑ^ and BD^−^grid-cell subgroups points in the same direction. Non-grid cells behave in an almost identical manner (Fig. 7*A*) as grid cells, see also Fig. 7-1.

Consistent with the hypothesis of one single group of bursty neurons, The physiological properties of individual cells could either be fixed or undergo plastic changes that move the biophysical cell parameters between the BD^ᴑ^ and BD^−^regions. In the ΔV_1_-ΔV_2_ space (Figs. 1C,7A), a transition from BD^ᴑ^ to BD^−^ corresponds to an increase in ΔV_1_ accompanied by a somewhat smaller decrease in ΔV_2_. Such a parameter change can be achieved through modifications of the AP-threshold, fAHP-minimum and/or DAP-maximum, as illustrated by the arrows in Fig. 1-1. Various ion channels have been implicated in DAP generation, from sodium and calcium channels (Alessi et a., 2016), to potassium (Eder et al. 1991) and HCN channels (Dickson et al., 2000), which also play a key role for slower grid-cell rhythms (Giocomo and Hasselmo, 2009). These channels could be regulated, e.g., by cholinergic stimulation, which has been shown to induce DAPs and after discharges in MEC-Layer-II neurons (Magistretti et al., 2004). Such modulations would have a direct impact on the precise temporal characteristics of bursting neurons.

Modulations of the biophysical parameters governing the afterpotentials might even occur at the time scale of single runs through the animal’s environment. Indeed, close inspection of individual membrane-potential traces suggests that BD^ᴑ^ cells do not generate a DAP after every AP; conversely, some action potentials of BD^−^ cells are followed by a DAP. One might even speculate that most bursty cells are capable of generating DAPs – slice experiments in rats suggest 85% of Layer-II stellate cells and 73% of Layer-II pyramids have DAPs (Canto and Witter, 2012) – but that this mechanism is under external control so as to switch cells between BD^ᴑ^ and BD^−^ behavior.

Remarkably, the ISI distributions of BD^−^ cells have ultra-sharp peaks, whose location varies only minimally within that group. Notably, the same short ISIs are elicited by the sparsely bursting neurons in Layer III (see also Mizuseki et al., 2009) and could be mediated by specific couplings between Layer-II BD^−^ cells and Layer-III SB neurons. The precise function of burst sequences in the 250-300 Hz regime remains an open question. Similarly, it is not obvious how cells with highly distinct firing characteristics can be orchestrated to create one joint grid-cell network (but see Pastoll et al., 2013), in which the SB, BD^−^ and BD^ᴑ^ cell classes have roughly the same grid score, spatial information and head direction score (Fig. 5*E-G*). With their high rate of bursts, BD^−^neurons might be ideally suited to drive other neurons in the network, whereas the DAPs of BD^ᴑ^ cells might trigger synaptic plasticity, similar to their function in CA3 pyramidal neurons (Mishra et al., 2016), and thus play a critical role for network reconfiguration when the animal learns about new environments (Krupic et al., 2018) or goals (Boccara et al., 2019).

Switching on the DAP mechanism (without interfering with the preceding fAHP) would then increase the probability of additional APs (Alessi et al., 2016) as well as provide a trace for the long-term potentiation of incoming synapses (Mishra et al., 2016). Once these synapses are strengthened and the DAP mechanism has been turned off again (or masked), the cell can fire precisely tuned bursts with short ISIs. These cell-intrinsic processes could be complemented by precisely wired and timed synaptic inputs (Varga et al., 2010; Couey et al., 2013; Pastoll et al., 2013; Buetfering et al., 2014; Fuchs et al., 2016; Schmidt et al., 2017; Winterer et al., 2017). Through short-term plasticity and integrative postsynaptic processes (Lisman, 1997; Izhikevich et al., 2003) such reorganization could result in a stronger influence on downstream neurons.

In contrast to what one might have expected, the strong dependence of DAPs on the neuron’s recent history (Alonso and Klink, 1993; Canto and Witter, 2012; Alessi, 2016) does not seem to translate into a spatial burst code. For example, one might have hypothesized that the DAP of a Layer-II stellate cell should be particularly large when the animal is moving into one of the cell’s firing fields, as this corresponds to raising the membrane potential from its previous out-of-field hyperpolarization. However, we could not find any signature for the ring-like burst-field structure expected in this scenario. In fact, we could not find *any* spatial dependencies despite vigorous search. This came as a surprise, given the role of burst firing for spatial coding in the hippocampus (Harris et al., 2001) or subiculum (Simonnet and Brecht, 2019). Similarly, spike doublets do not seem to play any special role for burst coding. Together, these findings suggest that grid-cell bursts are either not utilized for spatial coding, apart from their contribution to theta-phase precession (Hafting et al., 2008, Reifenstein et al., 2012), or that the spatial coding is masked by temporal fluctuations that are uncorrelated with spatial coordinates.

It is well known that after-spike potentials play a critical role in the control of AP firing patterns. For example, medium afterhyperpolarizations control theta-band clustering of action potentials in MEC stellate neurons (Fransen et al., 2004; Pastoll et al., 2013). Our study extends these and related findings to the 250-300 Hz range and provides a novel mechanistic explanation of burst firing of principal neurons in medial entorhinal cortex.

## Extended Data

**Figure 1-1.**
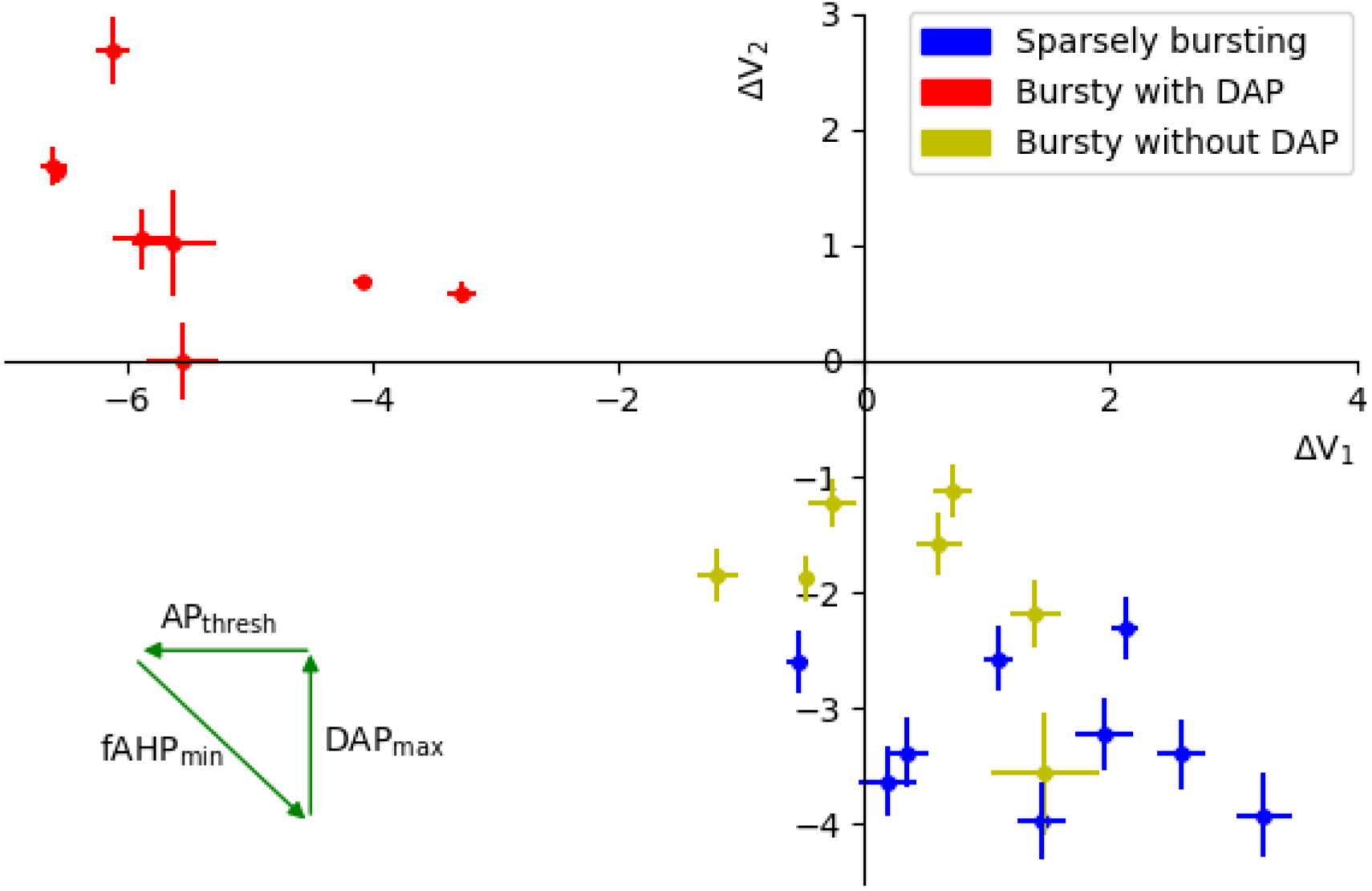
Stability of afterpotential parameters. To quantify the reliability of the parameters characterizing the spike afterpotentials, we carried out a bootstrapping analysis. Error bars indicate s.e.m. as obtained from 1000 samples and demonstrate that the fAHP and DAP parameters can be reliably estimated. The three arrows show how a cell’s position in the ΔV_1_-ΔV_2_ space changes when the respective parameter is increased.

**Figure 4-1.**
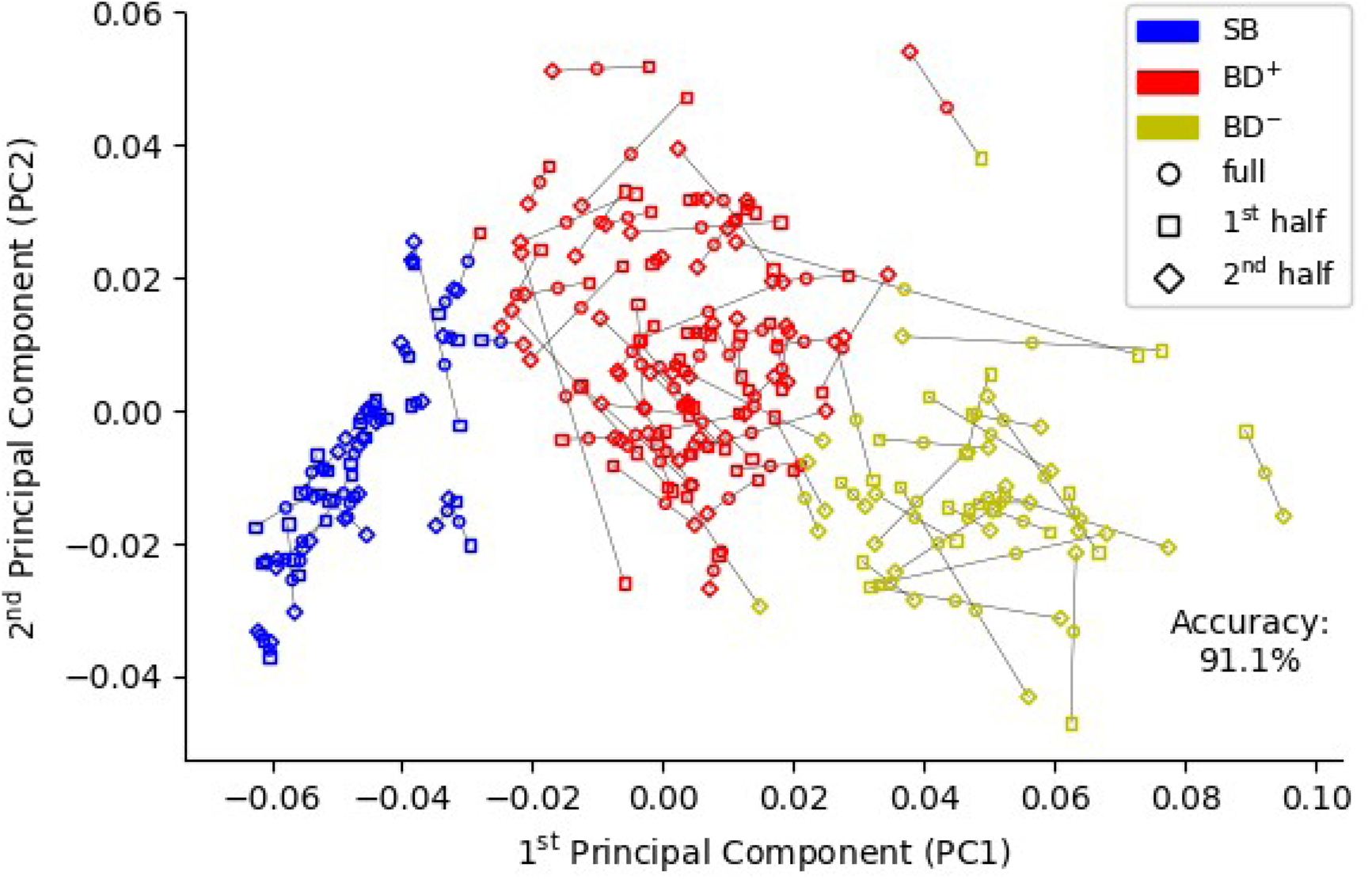
Robustness of cluster analysis. To test the robustness of the PCA-based class assignment of the grid-cell data from Latuske et al. (2015), we separately considered the first and second half of all spikes for each neuron. We then computed the autocorrelations within these two sets and projected the results into the PC space of the full grid-cell data. K-means clustering (k=3), carried out in the same way as had been done for the full data set, results in some neurons switching group identity. But only 8.9% of cells switch identity, when 1^st^ and 2^nd^ halves are compared, underscoring the robustness of the cluster analysis.

**Figure 6-1.**
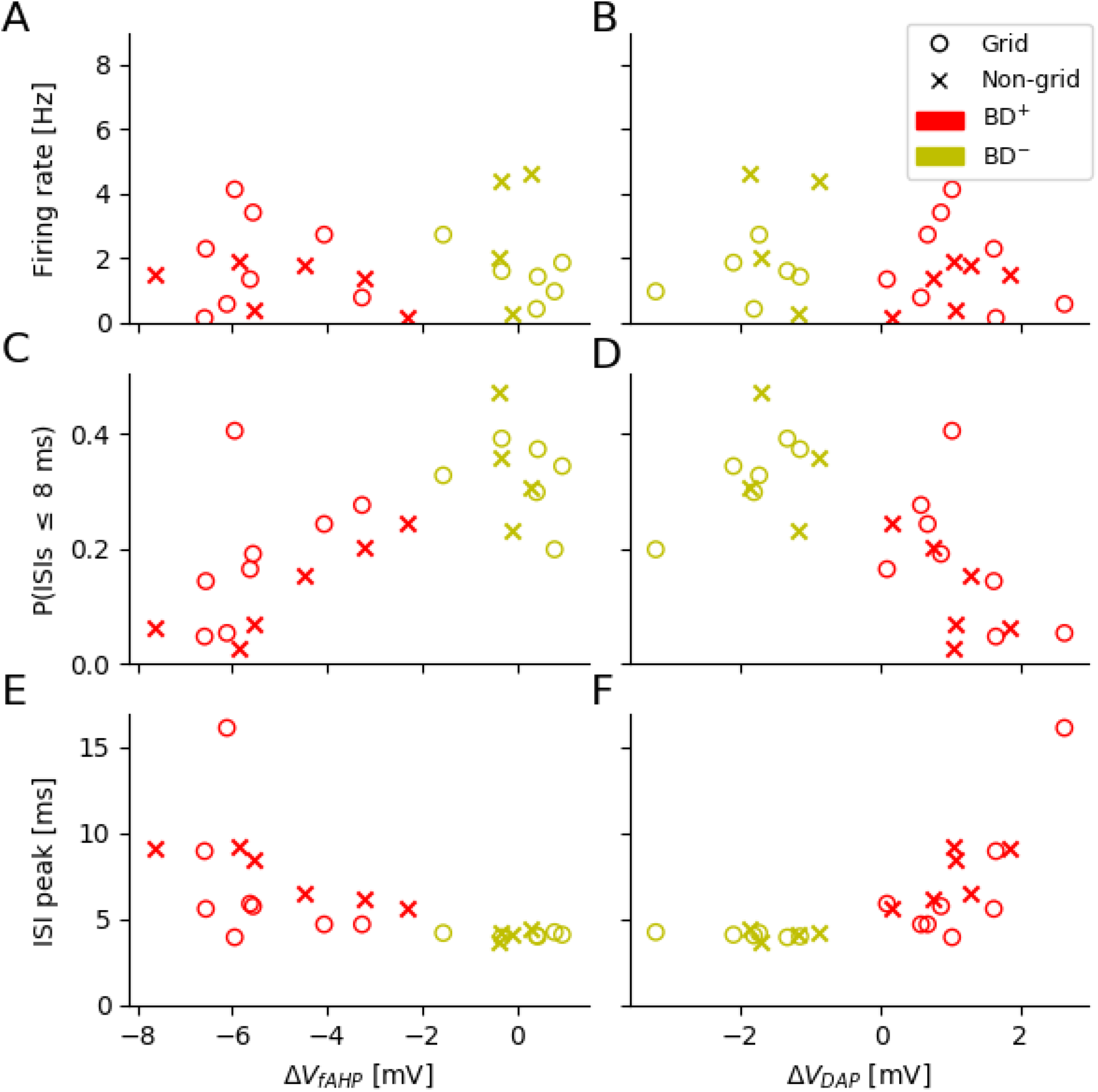
Cluster structure of all bursty principal cells recorded in virtual reality. Dependence of key spike-train parameters on fast afterhyperpolarization (ΔV_1_) and afterdepolarization (ΔV_2_). **A** and **B**, mean firing rates. **C** and **D**, fraction of ISIs below 8 ms (“burstiness”). **E** and **F**, location of ISI peak. There is no apparent difference to the grid-cell cluster structure shown in Fig. 6.

**Figure 7-1.**
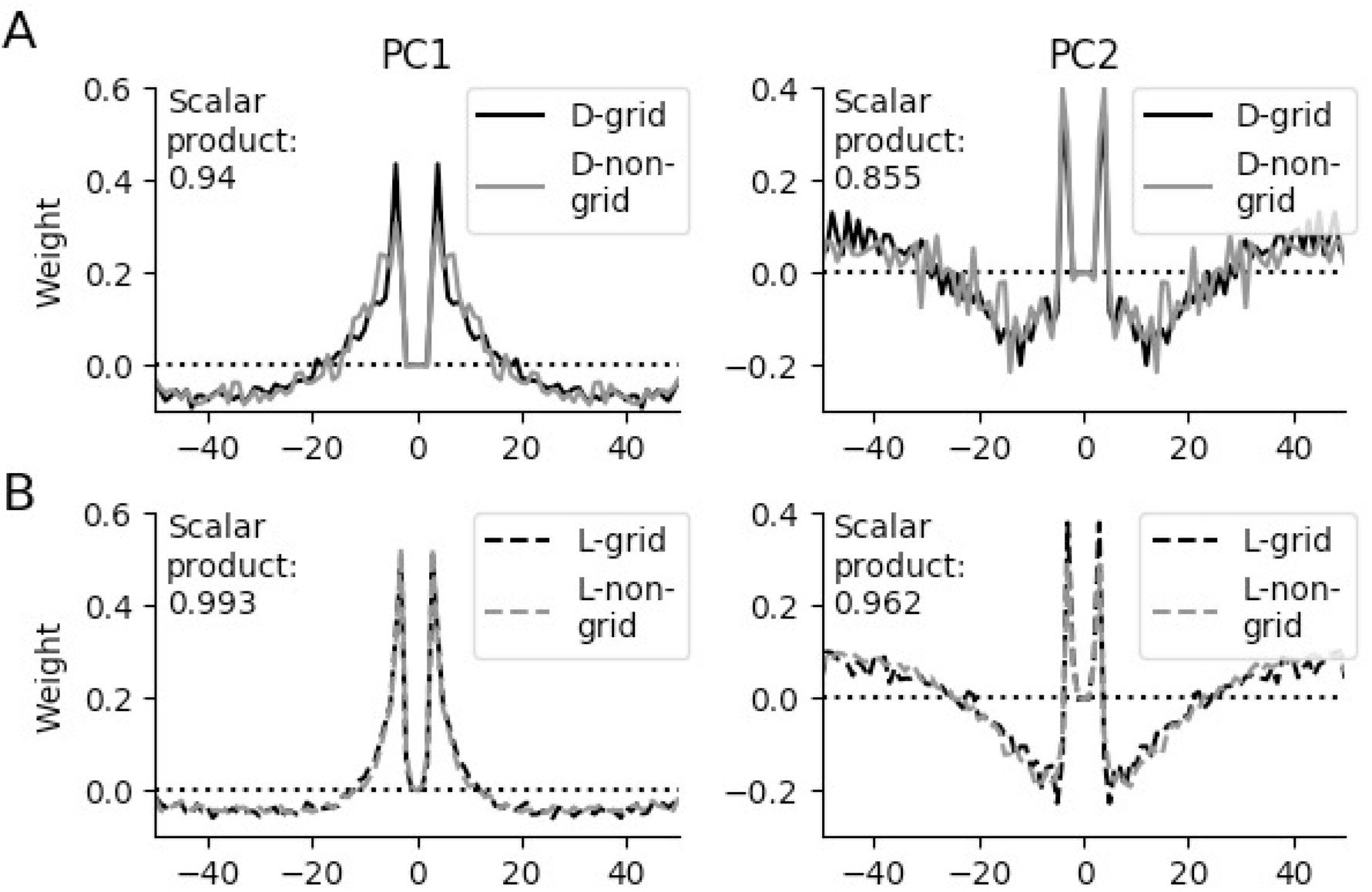
Principal components of the spike-time autocorrelations of grid cells and non-grid cells. **A**, First principal components (left panel) and second principal components (right panel) of the dataset from Domnisoru et al. (2013) for grid cells and non-grid cells. The similarity of the PC components is measured by the scalar product for time lags up to 50 ms. **B**, As in **A**, but now for the dataset from Latuske et al., (2015). The values of the scalar products are rather close to their maximal value of 1 and underscore the similarity of the grid-cell and non-grid-cell autocorrelations and their cluster structure.

